# Harsh Environments and “Fast” Human Life Histories: What Does the Theory Say?

**DOI:** 10.1101/014647

**Authors:** Ryan Baldini

## Abstract

A common belief among human life history researchers is that “harsher” environments - i.e., those with higher mortality rates and resource stress - select for “fast” life histories, i.e. earlier reproduction and faster senescence. I show that these “harsh environments, fast life histories” - or HEFLH - hypotheses are poorly supported by evolutionary theory. First, I use a simple model to show that effects of environmental harshness on life history evolution are incredibly diverse. In particular, small changes in basic but poorly understood variables - e.g., whether and how population density affects vital rates - can cause selection to favor very different life histories. Furthermore, I show that almost all life history theory used to justify HEFLH hypotheses is misapplied in the first place. The reason is that HEFLH hypotheses usually treat plastic responses to heterogeneous environmental conditions *within* a population, whereas the theory used to justify such hypotheses treat genetic responses to environmental changes across an entire population. Counter-intuitively, the predictions of the former do not generally apply to the latter: the optimal response to a harsh environment within a large heterogeneous environment is not necessarily the optimal strategy of a population uniformly inhabiting the same harsh environment. I discuss these theoretical results in light of the current state of empirical research.

## 1. Introduction

A popular hypothesis among evolutionary psychologists and anthropologists is the “harsh environments, fast life histories” - or HEFLH - hypothesis. This broadly states that harsh environmental conditions - those with high mortality rates and resource stress - select for greater investment in early reproduction at the expense of survival and later reproduction [1, 2, 3, 4, 5, 6]. Formal theoretical support rarely accompanies HEFLH hypotheses. The intuitive argument, however, is as follows: in high-mortality, resource-stressed environments, individuals have a smaller probability of surviving to any given age, including reproductive maturity. Those who die prior to or during reproductive ages pass fewer genes onto future generations, so selection should favor heritable strategies that shift energy toward early reproduction and faster senescence. Such thinking dates back at least fifty years to G.C. Williams’s [7] theory of senescence. These arguments are widely appealing: one paper [4] called the “high extrinsic mortality → early reproduction” result a “fundamental prediction” of life history theory. In fact, life history theorists have known for at least twenty years that this verbal argument is, in general, wrong [8, 9, 10], but this knowledge has not diffused into the anthropology and psychology literature.

Mechanistically, most HEFLH hypotheses invoke phenotypic plasticity, particularly as a response to early-life conditions [1, 2, 11, 12] (but see [13] for a genetic view of life history strategies in humans). That is, various stressors in infancy and childhood are thought to predict resource stress and mortality risk throughout life. Children therefore adaptively respond by speeding up their reproductive schedule - e.g., by maturing early, reproducing early and often, and devoting less energy to childcare and longevity [1, 2, 4].

The two purposes of this paper are to show that

1. Evolutionary theory actually does not, in general, predict that harsh environments favor fast life histories. Harsh environments, however defined, have a diversity of effects that depend on various mechanistic assumptions.
2. The life history theory cited in support of HELFLH hypotheses are inappropriate when applied to cases of phenotypic plasticity.

Regarding point (1): I build a simple model to show that the effects of harsh environments on life history evolution are diverse. Specifically, the effects depend on a variety of poorly understood biological variables, including the exact way in which harshness affects survival and fertility at different ages, and how population density affects population growth. Successful application of life history theory to humans therefore requires more detailed knowledge of human demography and physiology. Such details are rarely, if ever, mentioned in the HEFLH literature.

Regarding point (2): almost all life history models cited by HEFLH hypotheses (e.g., [14, 15]) have focused on the evolutionary response when an entire population is subjected to a change in environmental conditions. The majority of HELFH hypotheses, however, involve plastic responses to environmental conditions that vary *within* a population. By applying the former theory to the latter problem, researchers implicitly assume that an individual inhabiting a harsh subenvironment within a heterogeneous population will be selected to behave the same life history as an individual in a population uniformly inhabiting the same environment. I show that this is often false, implying that the theory they cite is inappropriate in the first place. Future efforts to evaluate HEFLH hypotheses will need to produce models that explicitly account for phenotypic plasticity.

The following two sections treat these points in turn. I then discuss these findings in light of what is currently known about life history theory and human reproductive flexibility.

## 2. Harsh Environments have Diverse Effects on Life History Evolution

Using a simple model, I derive a few cases in which increased environmental harshness does not favor fast life histories. From a hypothesis testing framework, this proves that HEFLH hypotheses are not a necessary consequence of life history theory; that hypothesis is quickly rejected. More importantly, however, I demonstrate just how sensitive life history evolution is to very subtle changes in basic biological assumptions. Taking one simple model, I show that slight changes in (1) the precise way in which harsh environments affect survival and fertility and (2) the nature of population regulation can lead to opposite outcomes of life history evolution. Life history theory can do little for us until we gain a better understanding of these basic demographic details for human populations.

Consider an asexual, age-structured population. Throughout their lives, individuals devote a proportion *q* of available energy to mortality reduction, e.g., by improving immune function and DNA repair. The remaining proportion 1 − *q* is devoted either to growth (prior to maturity) or reproduction (after maturity). Individuals with a larger *q* therefore live longer, at the expense of reproductive output. Individuals grow until age *a*, at which point they begin reproduction. Reproductive output in adulthood is proportional to one’s size; therefore, increasing *a* implies a later age at first reproduction, but a higher fertility rate once maturity is reached. I assume that *q* and *a* are simultaneously under evolution; i.e., the population exhibits genetic variation in both of these variables, but no other (which are introduced below). In reality, of course, genetic variation can affect any parameter of the model, but considering just these two will be enough to prove the point.

To complete the model, we must specify whether and how population density affects survival and reproduction, and write these as mathematical functions. I treat two cases in turn: (1) density-independent (exponential) population growth, and (2) density-dependent fertility across all ages. One can show that case (1) also applies to density-dependent mortality for all ages, and that case (2) also applies to density-dependent infant survival, so these are more widely applicable than they appear. Still, these are just two of many possible forms of population regulation; the diversity of outcomes would surely be greater still if a wider variety of assumptions were explored.

### 2.1. Density-independent population growth

Suppose that the age-specific survival and fertility rates are not affected by population size. Specifically, let the function *l* be the age-dependent survival function, such that *l*(*x*) is the probability of surviving to age *x*. Let the function *m* be the age-dependent fertility function, such that *m*(*x*) is the expected number of offspring to an individual of age *x*. I assume these take the following form.

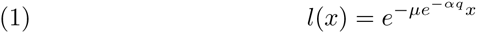

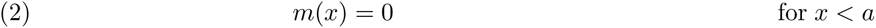

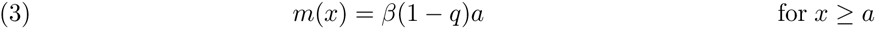

**Table 1.** The effect of increased harshness on the optimal values 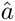 and 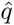. Each row describes a different kind of increased environmental harshness, as described in section 2.1. The symbol ↑ means that increased harshness favors “slower” life histories, and vice versa for ↓. The symbol “−” signifies no effect. The double arrow symbol ↑↓ means that the effect can go either direction, depending on other parameter values.

|  |  | Population Regulation |  |
| --- | --- | --- | --- |
|  |  | Density independence (2.1) | Density-dependent fertility (2.2) |
| Harshness | $\mu \uparrow$ | $\hat{a} \uparrow, \hat{q} \uparrow$ | $\hat{a} \downarrow, \hat{q} -$ |
| | $\alpha \downarrow$ | $\hat{a} \uparrow, \hat{q} \uparrow$ | $\hat{a} \downarrow, \hat{q} \downarrow$ |
| | $\beta \downarrow$ | $\hat{a} \uparrow\downarrow, \hat{q} \uparrow\downarrow$ | $\hat{a} -, \hat{q} -$ |

Equation (1) says that the probability of surviving decreases exponentially as one gets older, due to a constant death rate of *μe*^−*αq*^. This means that *μ* is the death rate if no energy is invested in survival (i.e., *q* = 0), and that the death rate decreases exponentially at rate *α* as individual invests more in survival. Equations (2) and (3) imply that reproduction is 0 prior to maturity at age *a*. The eventual reproductive rate is proportional to the amount of time spent growing as a juvenile (*a*) and the amount of energy invested in growth and reproduction (1− *q*). *β* is an exogenous parameter that determines how easily energy is converted to growth and reproduction.

For any set of parameters *μ*, *α*, and *β*, it is possible to find the optimal values in *a* and *q*. Denote these optimal values by 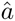 and 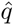. We now ask: *what is the effect of increased environmental harshness on the optimal values 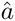 and 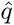?* To answer this question first requires that we define “environmental harshness” precisely. In this model, we can increase environmental harshness in three ways. First, we can increase the extrinsic mortality rate, i.e, *μ* ↑. Second, we can decrease the effectiveness of investments in mortality reduction, i.e., *α* ↓. This decrease in *α* means that investments in survival are less effective; one must invest more to achieve the same rate of survival as in a less harsh environment. Third, we can decrease the effectiveness of investments in growth and fertility, i.e., *β* ↓. A decreased *β* means that one must spend a longer time growing to reach the same fertility rate as in a less harsh environment. I consider all three effects, in turn.

I show how to analyze the effects of these changes in the appendix. Table 1 summarizes the results. First, 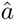 and 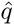 both increase with *μ*; contrary to HEFLH predictions, increasing the extrinsic mortality rate favors later reproduction and less reproductive effort overall. Second, 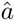 and 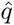 both increase as *α* increases. This means that as mortality reduction gets more “expensive” (*α* ↓), selection again favors a slower life history strategy, with 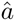 and 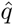 both increasing. Third, changing *β* can either increase or decrease 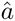 and 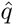, depending on the values of other parameters. The general trend here is that harsh environments appear to favor *slower* life histories, although in one case (*β*) the effect is ambiguous. The latter fact shows that the precise definition of “harshness” is important, as different kinds of harshness can have different effects.

Why does this happen? Explaining these results is very difficult. In general, a change in any extrinsic parameter will have a direct selective affect on *a* and *q*. It will also change *r*, the population growth rate, which is also a central parameter in life history evolution. Finally, the selection gradients for *a* and *q* depend on each other (see the appendix), which means that evolution in one will alter selection on the other - i.e., they *coevolve*. The interaction of all these moving parts makes “intuitive” understanding an unrealistic goal. Fortunately, intuition is not necessary: math does the work for us.

### 2.2. Density-dependent fertility

The results of life history evolution depend strongly on how population density affects population growth. To demonstrate this, we now take the same model as above, but suppose that fertility at all ages is depressed by population density. In particular, assume that fertility decreases exponentially with total population size, at rate *D*. Then

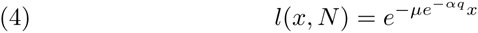

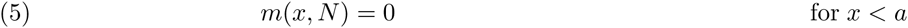

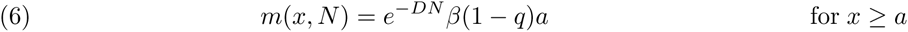

The remaining parameters have the same meaning as in section 2.1.

How does this innocuous demographic change affect evolution in *a* and *q*? Fortunately, in this case, it is possible to solve for the optimal values 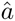 and 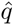, algebraically (see appendix). They are

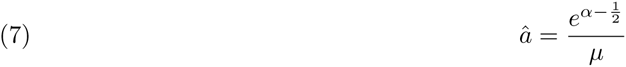

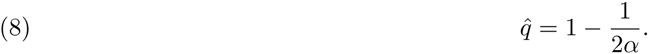

We can deduce all important results rather quickly. First, note that *β* is absent: the cost of growth and reproduction has no effect on life history evolution in this case. Second, *μ*, the extrinsic mortality rate, has no effect on 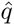, the optimal allocation to survival investment, nor therefore any effect on the rate of senescence - in contrast to the HEFLH hypotheses. On the other hand, 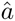, the optimal age at reproduction, does evolve downward as the extrinsic rate of mortality increases. This is consistent with many HEFLH hypotheses, but opposite to the density-independent case analyzed just above. Finally, any decrease in *α*, which measures the effectiveness of survival investment, will select for earlier age at first birth and lesser investment in survival. We conclude that simply imposing a particular form of density dependence has altered *every* environmental effect (Table 1).

## 3. Plastic Responses to Heterogeneous Environments: Classic Predictions may not Apply

There is a second, and perhaps more fundamental, problem with HEFLH hypotheses. HEFLH hypotheses usually treat *plastic responses* to harsh conditions *within* a heterogeneous population. In other words, they hypothesize that an individual born into harsh conditions will adaptively respond by employing a fast life history [1, 2, 4, 11, 12]. Those who are born into more fortunate conditions are expected to employ a slow life history. In most cases, these are not assumed to be due to genetic differentiation between people in different conditions, but to plastic response. In contrast, the theory cited to justify these hypotheses (e.g., [14, 15]) assumes that an *entire* population is genetically evolving toward a *uniformly* changed environment. The models analyzed above did this too: we imagined that an entire population is thrown into a harsh environment, and asked how life history evolved in response, across many generations.

Intuition suggests that this should not be a problem. We might suspect that the optimal plastic response to a harsh “sub-environment,” within a broader heterogeneous populations, would be exactly the optimal fixed behavior of a population that uniformly inhabits the same conditions. This is tacitly assumed when classic life history theory is used to justify HEFLH hypotheses. In fact, this intuition is wrong. This means that the application of most life history theory to HEFLH hypotheses is fundamentally inappropriate.

I derive this surprising result in the appendix. It can be understood as follows. Suppose first that an asexual population uniformly inhabits a particular environment. Suppose the population is composed solely of a single genetic variant, which yields (using the notation of section 2) survival function *l* and fertility function *m*. These cause the population to grow at a constant rate *r*. Suppose that a rare variant arises which yields slightly changed functions *l*′ and *m*′. Standard population genetic methods [9] show that the new variant will invade and replace the common one only if

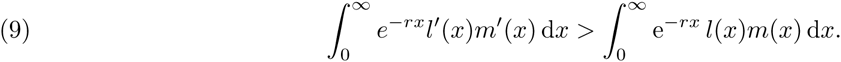

It is possible to show (see [9]) that this equivalent to the condition

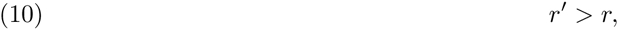

where *r*′ is the growth rate of a population consisting only of the invading variant. That is, the population with the faster growth rate eventually displaces the other.

Now consider a population that inhabits a heterogeneous environment. For simplicity, suppose that there are two sub-environments, called *A* and *B*. For example, these could be represented by different parameter values in the models discussed in section 2 - perhaps *B* has a higher extrinsic mortality rate *μ*. With probability *k*, an individual is born into *A*, and remains there; otherwise, she inhabits *B*. Further, suppose that individuals can accurately detect which sub-environment they inhabit, and can respond to each by employing different life history strategies. The plastic response is encoded in the genotype: it specifies what to do in each environment. For the resident genotype, inhabiting sub-environment *A* invokes survival and fertility functions *l_A_* and *m_A_*; *B* invokes *l_B_* and *m_B_*. This population will grow at a rate 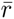, which lies between the would-be growth rate of a population solely inhabiting *A* (*r_A_*) and another inhabiting *B* (*r_B_*). Now suppose a new variant arises that only alters the plastic response to sub-environment *A*, yielding functions 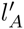 and 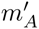. This is a plasticity mutation: it affects behavior only in a particular environmental condition. The appendix shows that this mutant invades and replaces the other only if

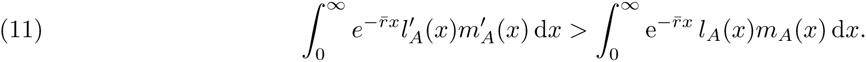

It turns out that this is *not* equivalent to the condition 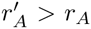, which would be the invasion condition if the entire population inhabited environment *A* (as in equation (10)). The reason, mathematically, is that 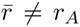; the presence of other sub-environments alters the total population growth rate, which alters the “discount rate” of later reproduction for all sub-environments^1^.

The same procedure for a density-dependent population shows that a plasticity mutation affecting life history in environment *A* invades only if

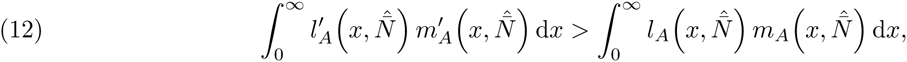

where 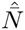 is the equilibrium population size of the entire population, which will lie somewhere between 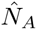 and 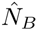. Equation (12) is, again, not equivalent to the condition 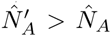, which would be the invasion condition if the entire population inhabited environment *A* [9].

For a concrete example, let us take the density-independent model from section 2.1, and put it in a variable environment. For simplicity, assume that only *μ* varies between sub-environments, yielding two parameters *μ_A_* and *μ_B_*. Let *B* be the harsher environment, so *μ_B_* > *μ_A_*. Assume that an individual has equal probability of being birthed into each environment, i.e., *k* = 0.5.

We ask: what is the optimal plastic response in each environment? Answering this question in general is difficult, but one can find the optimal strategies numerically, for any given set of parameters (see appendix). Figure 1 shows optimal responses for a range of *μ_B_*, when *μ_A_* is held at 1 (see Figure 1 caption for other parameter values). A surprising but important fact appears: a change in *μ_B_*, the harshness of sub-environment *B*, changes the optimal life history strategies (in *a* and *q*) for *both sub-environments A* and *B*. This is because increasing mortality in *B* depresses the entire population’s growth rate, which affects evolution in both sub-environments.

**Figure 1.**
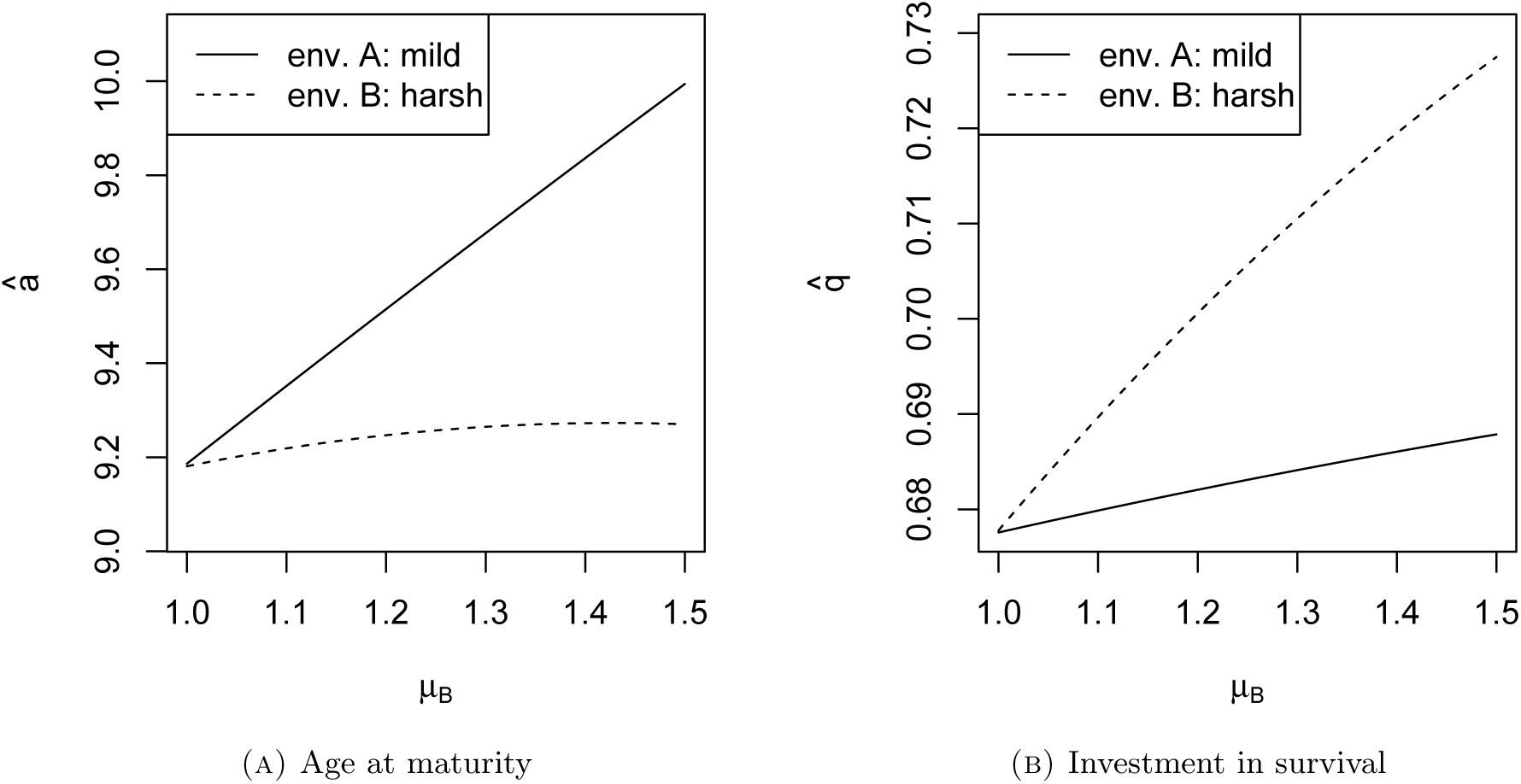
Optimal plastic responses for *a* and *q*. *μ_A_* = 1, *α* = 5, *β* = 0.1, *k* = 0.5. The plotted range shows *μ_B_* ≥ *μ_A_*, so *B* is the harsher environment.

Figure 1 also shows how the results of classic life history theory can be misleading for plastic life history responses. A straightforward application of section 2.1 would lead us to predict that individuals in sub-environment *B* should mature later and invest more in survival than in *A* (see Table 1). Figure 1 shows that this is false: although those in *B* do invest more in survival, they actually mature earlier than those in *A*. This demonstrates that one cannot simply take the results of classic life history models and apply them to cases of life history plasticity; one must model the evolution of plasticity explicitly.

Finally, this example shows that the “fast-slow” life history dichotomy can fail. Individuals in the harsh sub-environment *B* reach reproductive maturity earlier those in *A*, which is usually considered a fast life history trait. But they also invest less in reproduction overall, and senesce more slowly than those in *A* - a “slow” life history trait. Are individuals in environment *B* fast or slow? There appears to be no precise, quantitative measure of life history speed, so this question ultimately cannot be answered. In any case, there is little reason to expect that all life history traits will evolve in concert along a single, intuitive dimension.

## 4. Discussion

The main message of this paper - that harsh environments can favor any life history strategy depending on the physiological and demographic details - is frustrating. Are there no generalities that we can apply without knowing, for example, the precise mechanisms of density dependence in ancestral human populations? Theorists have suggested some possibilities, but in fact none have turned out to be sufficiently general. Abrams [8] investigated the effect of extrinsic mortality under a variety of density-dependent scenarios. The results he provides are attractive but may not hold when mutations have complex effects on vital rates. For example, he claims that, under density independence, “if the extrinsic survival probability… is lowered by a factor *k* (*k* < 1), then [there is] no change in the evolutionarily favored rate of senescence” ([8], p.880). The density-independent case of section 2.1 (see Table 1) shows that this false: a proportional increase in the mortality at all ages (*μ* ↑) selected for slower senescence 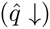. Similarly, Charnov’s “invariant” life history rules [15, 16] are derived under the assumption of population stationarity resulting from density-dependent juvenile recruitment (equivalent to the model of section 2.2). They do not necessarily hold under other forms of density dependence, e.g., density-dependent adult mortality.

Rather than searching for theoretical generalities, a more fruitful line of research would seek to understand the essential mechanisms highlighted in this paper. How precisely do reproduction and survival trade off at different ages? Which ages do harsh environments most affect? Does harshness affect survival, fertility, or both? What demographic mechanisms have regulated human population growth in the past? How does density dependence operate? Each of these issues plays a central role in life history theory; we won’t be able to make strong evolutionary predictions until we’ve answered them.

None of the theoretical work in this paper refutes the empirical findings discovered by various tests of HEFLH hypotheses. It does, however, cast doubt on the evolutionary justification and interpretation of such findings. Many studies have indeed found that stressful infant or childhood environments - as measured by, e.g., infant birth weight, familial warmth or positivity, father absence - are correlated with earlier sexual maturation and reproduction in humans (see reviews in [17, 12, 18]). A primary difficulty with such correlation studies is, of course, demonstrating causation [6]. It may be, for example, that people who inhabit such environments are genetically inclined to have “fast” life histories in the first place. Distinguishing correlated genetic and environmental effects is difficult, especially since stressful childhood conditions may be largely shared among siblings. One children-of-twins study suggests that genetic variation may account for much of the observed negative correlation between daughter’s menarcheal age and stepfather presence [19]. Similarly, sisters reared together, but differentially exposed to paternal dysfunction and parental separation, did not significantly vary in menarcheal age - although there was a significant interaction between the two [20]. Perhaps the most straightforward test would be to measure the effect of childhood stressors on various life history characteristics among monozygotic twins separated at birth; this would provide perfect genetic control, albeit at the expense of representation. In any case, it is clear that genetic and plastic hypotheses are not mutually exclusive.

## Appendix A. Derivations for Section 2

### A.1. Section 2.1: Density independence

For density independence, a mutant value of *a* will invade only if it has a faster growth rate than that of the resident strategy. *r* is found by solving the equation

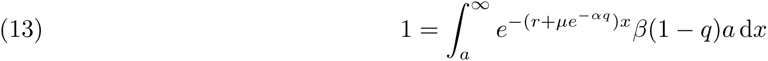

Therefore, we can deduce the results of selection by solving the above equation for *r* and differentiating with respect to *a*; a positive derivative implies that *a* will evolve upward. The same applies for *q*. This procedure assumes that evolution proceeds primarily by the substitution of infrequent mutations.

Solving equation (13) yields

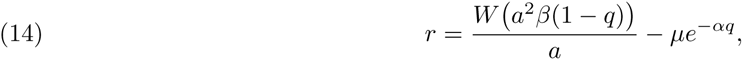

where *W* is the product-log function: *W* (*x*) is the number that solves the equation *x* = *W* (*x*)*e*^*W* (*x*)^. Proceeding with differentiation, we first find that

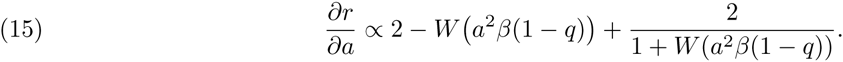

When *a* = 0, the product-log term is 0 and the above gradient is positive. It follows that *a* always evolves upward from 0. Similarly, as *a* → ∞, the product-log term → ∞, and therefore the above gradient → −∞. This implies that the optimal value of *a* is indeed positive and finite. Setting (15) to 0 shows that, at evolutionary equilibrium, this simplifies to *W* (*a*^2^*β*(1 − *q*)) = 1, or

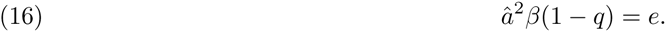

Differentiating (14) with respect to *q*, we find

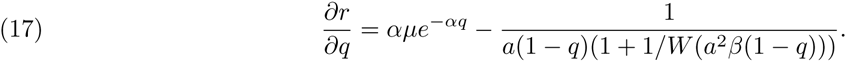

When *q* = 1, equation (17) reveals that *r* < 0, so *q* = 1 cannot be optimal in any biological realistic context. When *q* = 0, (17) evaluates to

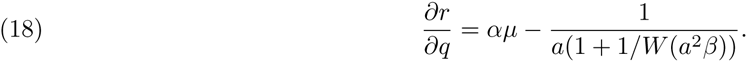

If the right-hand side of (18) is less than 0, then the optimal value of *q* is 0; the population will evolve such that individuals invest no energy in mortality reduction. If not, then *q* will evolve to some intermediate optimum, which is found by setting (17) equal to 0. Doing this, and substituting (16), yields the equilibrium condition

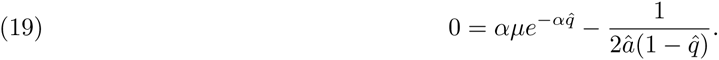

The optimal values 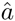 and 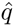 are found by simultaneous solution of (16) and (19) for *a* and *q*. This system cannot be solved algebraically, but numerical solutions can be found for any set of parameters, e.g. via the R package *rootSolve*.

We can characterize the effect of changing the parameters *μ*, *α*, and *β* on 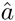 and 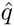. To do this, we express 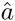 and 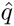 as functions of one of these parameters, e.g., 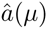 and 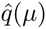. We substitute these functions into equations (16) and (19), and implicitly differentiate both with respect to the parameter of interest. This yields to equations involving the unknown optima and their derivatives, i.e., 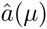, 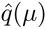, 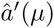, and 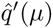, where primes here denote derivatives. We can then solve for the derivatives, which tell us how optima change with changing environmental parameters. Although we do not know the optimal values in algebraic terms, we do know that 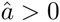 and 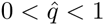, which, fortunately, allows us to determine the sign of some derivatives.

Using *Mathematica* to take partial derivatives and solve the resulting system of equations, I find (ignoring positive factors, and dropping the arguments of 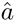 and 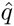 on the right-hand side)

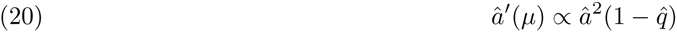

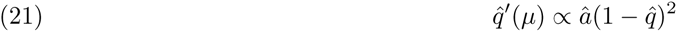

Both expressions are necessarily positive whenever the equilibrium exists. Therefore, increasing *μ* selects for larger *a* and *q*.

Repeating for *α*, we have

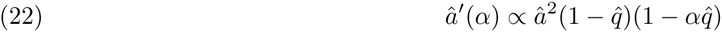

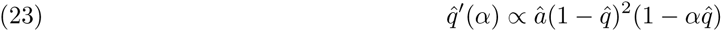

signs of both expressions are determined by 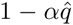. This suggests that the effect *α* depends on value of other variables; sometimes the effect is positive, and sometimes negative. This is confirmed by exploring parameter values with *rootSolve* in R.

Finally for *β*, we have

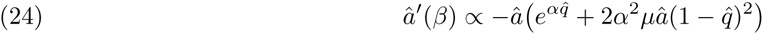

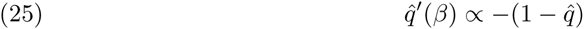

Both expressions are negative. Therefore, decreasing *β* selects for larger *a* and *q*.

### A.2. Section 2.2: Density-dependent fertility

I assume a genetically monomorphic population approaches an equilibrium population size 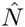. This is found by solving the equation

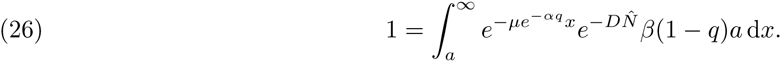

Charlesworth [9] shows that selection maximizes 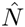 in such cases. We proceed as before, but replacing *r* with *N*. Solving equation (26) for 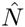 yields

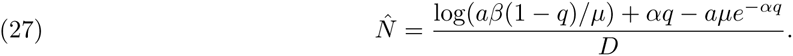

For *a*, we find

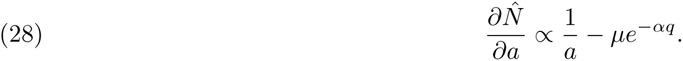

The above gradient goes to ∞ as *a* → 0, so *a* = 0 is not evolutionarily stable. Similarly, as *a* → ∞, the above gradient goes negative. We conclude that there is an intermediate optimum in *a*.

For *q*, we have

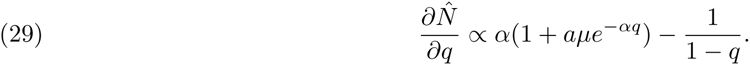

As *q* → 1, the above gradient goes to −∞, so *q* = 1 is not stable. If *q* = 0, then we find the condition for the increase in *q* to be

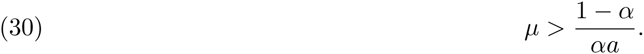

Thus, if equation (30) does not hold, then the optimal value of *q* is zero. Otherwise, *q* evolves upward from 0 to some intermediate optimum. In this case, we find the optimum by setting (28) and (29) equal to 0 and solving for *a* and *q*. Doing so yields equations (7) and (8) in the text.

## Appendix B. Derivations for Section 3

I treat only an asexual reproducing population, for simplicity; the same results can be extended as in [9]. I will derive the conditions for the invasion of a rare mutant life history strategy, which will generally be denoted by primes (′), as in section 3. Let the frequency of this rare allele among newborns at time *t* be *p*(*t*). Let *N*(*x*, *t*) be the total number of individuals in the population of age *x* in the population at time *t*. Working in discrete time, we have

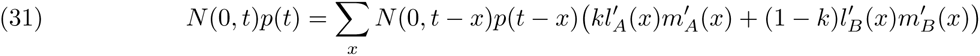

The quantity on the left is the total number of newborns at time *t* that are of the mutant strategy. This is equal to the sum of all mutant adults times their mean fertility - which is 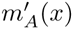 for individuals of age *x* in environment *A* and 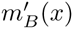 for those in *B*. For any age class *x*, the total number of mutants is equal to the number born at time *t* − *x* (*N*(0, *t* − *x*)*p*(*t* − *x*)) times the proportion who survived. This is 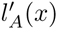 for those in environment *A*, etc. Dividing both sides by *N*(0, *t*) yields

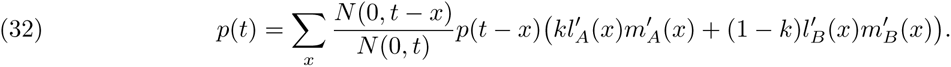

The ratio of population sizes, 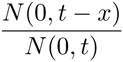, will usually rapidly converge on the value *e*^−*rx*^, where *r* is the growth rate of the resident strategy, i.e. the real solution to

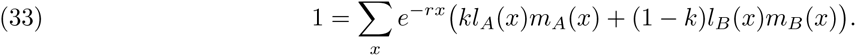

There will, in general, be a deviation from *r* due to the presence of the novel rare allele, but this deviation is negligible for a new, rare mutant. We then have

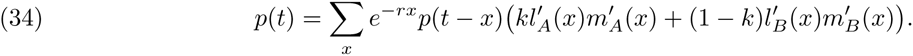

These equations are analogous to the dynamical equations in Charlesworth’s analysis of age-structured natural selection ([9], ch. 4). It follows that all the properties that apply there also apply here. In particular, the rare variant only invades if

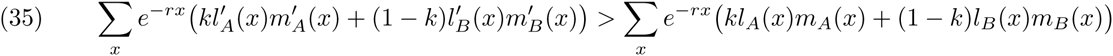

If a rare mutant only affect the survival and fertility function in one sub-environment - say, *A* - then (35) simplifies to equation (11) in the text (allowing a switch to continous time).

Following Charlesworth ([9], ch. 4), one can derive an analogous result under density dependence. We find that a new strategy invades only if

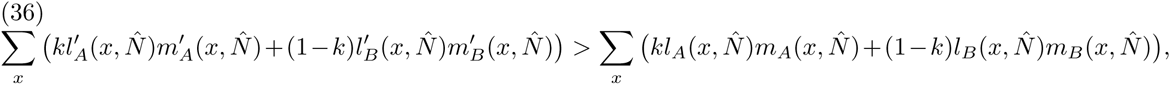

where 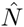 is the equilibrium population size of the resident strategy. If the resident strategy only affects the survival and fertility functions in one sub-environment - say, *A* - then (36) simplifies to equation (12) in the text (allowing a switch to continuous time).

Putting the model of section 2.1 into a two sub-environment scenario yields four evolving parameters: *a_A_*, *a_B_*, *q_A_*, and *q_B_*. The optimal life history strategy is found by maximizing *r*, subject to the constraint

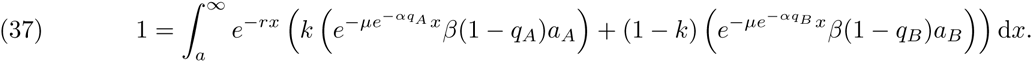

For Figure 1, I solved (37) for *r* using the function *uniroot,* in *R*. This can then be maximized with respect to the four evolving parameters using *optim*.

## Footnotes

1 The discount rate is found in the term *e*^−*r*^*x*; of equations (9) and (11). This term implies that later reproduction contributes less to fitness in growing populations whenever *r* > 0 [9]

## References

[1] Jay Belsky, Laurence Steinberg, and Patricia Draper. Childhood experience, interpersonal development, and reproductive strategy: An evolutionary theory of socialization. Child development, 62(4):647–670, 1991.

[2] James S Chisholm, Peter T Ellison, Jeremy Evans, PC Lee, Leslie Sue Lieberman, Zdenek Pavlik, Alan S Ryan, Elizabeth M Salter, William A Stini, and Carol M Worthman. Death, hope, and sex: Life-history theory and the development of reproductive strategies [and comments and reply]. Current anthropology, pages 1–24, 1993.

[3] Marco Del Giudice. Sex, attachment, and the development of reproductive strategies. Behavioral and Brain Sciences, 32(01):1–21, 2009.

[4] Bruce J Ellis, Aurelio José Figueredo, Barbara H Brumbach, and Gabriel L Schlomer. Fundamental dimensions of environmental risk. Human Nature, 20(2):204–268, 2009.

[5] Vladas Griskevicius, Andrew W Delton, Theresa E Robertson, and Joshua M Tybur. Environmental contingency in life history strategies: the influence of mortality and socioeconomic status on reproductive timing. Journal of personality and social psychology, 100(2):241, 2011.

[6] Daniel Nettle. Flexibility in reproductive timing in human females: integrating ultimate and proximate explanations. Philosophical Transactions of the Royal Society B: Biological Sciences, 366(1563):357–365, 2011.

[7] George C Williams. Pleiotropy, natural selection, and the evolution of senescence. Evolution, 11.

[8] Peter A Abrams. Does increased mortality favor the evolution of more rapid senescence? Evolution, pages 877–887, 1993.

[9] Brian Charlesworth. Evolution in age-structured populations. Cambridge University Press Cambridge, 1994.

[10] Hal Caswell. Extrinsic mortality and the evolution of senescence. Trends in ecology & evolution, 22(4):173–174, 2007.

[11] Tamas Bereczkei and Andras Csanaky. Stressful family environment, mortality, and child socialisation: Life-history strategies among adolescents and adults from unfavourable social circumstances. International Journal of Behavioral Development, 25(6):501–508, 2001.

[12] Daniel Nettle, David A Coall, and Thomas E Dickins. Early-life conditions and age at first pregnancy in british women. Proceedings of the Royal Society B: Biological Sciences, page rspb20101726, 2010.

[13] Aurelio Jose Figueredo, Geneva Vasquez, Barbara Hagenah Brumbach, and Stephanie MR Schneider. The heritability of life history strategy: The k-factor, covitality, and personality. Biodemography and Social Biology, 51(3-4):121–143, 2004.

[14] Stephen C Stearns. The Evolution of Life Histories, volume 249. Oxford University Press Oxford, 1992.

[15] Eric L Charnov. Life history invariants: some explorations of symmetry in evolutionary ecology, volume 6. Oxford University Press Oxford, 1993.

[16] Kristen Hawkes, James F OConnell, NG Blurton Jones, Helen Alvarez, and Eric L Charnov. Grandmothering, menopause, and the evolution of human life histories. Proceedings of the National Academy of Sciences, 95(3):1336–1339, 1998.

[17] Bruce J Ellis. Timing of pubertal maturation in girls: an integrated life history approach. Psychological bulletin, 130(6):920, 2004.

[18] Jay Belsky. The development of human reproductive strategies: Progress and prospects. Current Directions in Psychological Science, 21(5):310–316, 2012.

[19] Jane Mendle, Eric Turkheimer, Brian M D’Onofrio, Stacy K Lynch, Robert E Emery, Wendy S Slutske, and Nicholas G Martin. Family structure and age at menarche: a children-of-twins approach. Developmental psychology, 42(3):533, 2006.

[20] Jacqueline M Tither and Bruce J Ellis. Impact of fathers on daughters’ age at menarche: a genetically and environmentally controlled sibling study. Developmental psychology, 44(5):1409, 2008.

